# Palidis: fast discovery of novel insertion sequences

**DOI:** 10.1101/2022.06.27.497710

**Authors:** Victoria R. Carr, Solon P. Pissis, Peter Mullany, Saeed Shoaie, David Gomez-Cabrero, David L. Moyes

## Abstract

The diversity of microbial insertion sequences, crucial mobile genetic elements in generating diversity in microbial genomes, needs to be better represented in current microbial databases. Identification of these sequences in microbiome communities presents some significant problems that have led to their underrepresentation. Here, we present a bioinformatics pipeline called Palidis that recognises insertion sequences in metagenomic sequence data rapidly by identifying inverted terminal repeat regions from mixed microbial community genomes. Applying Palidis to 264 human metagenomes identifies 879 unique insertion sequences, with 519 being novel and not previously characterised. Querying this catalogue against a large database of isolate genomes reveals evidence of horizontal gene transfer events across bacterial classes. We will continue to apply this tool more widely, building the Insertion Sequence Catalogue, a valuable resource for researchers wishing to query their microbial genomes for insertion sequences.

**Data Summary:**

1. Palidis is available here: github.com/blue-moon22/palidis
2. The Insertion Sequence Catalogue is available to download here: https://github.com/blue-moon22/ISC
3. The raw reads from the Human Microbiome Project can be retrieved using the download links provided in Supplementary Data 1
4. The analysis for this paper is available here: github.com/blue-moon22/palidis_paper_analysis
5. The output of Palidis that was run on these reads is available in Supplementary Data 2

**Impact Statement:** Insertion sequences are a class of transposable element that play an important role in the dissemination of antimicrobial resistance genes. However, it is challenging to completely characterise the transmission dynamics of insertion sequences and their precise contribution to the spread of antimicrobial resistance. The main reasons for this are that it is impossible to identify all insertion sequences based on limited reference databases and that *de novo* computational methods are ill-equipped to make fast or accurate predictions based on incomplete genomic assemblies. Palidis generates a larger, more comprehensive catalogue of insertion sequences based on a fast algorithm harnessing genomic diversity in mixed microbial communities. This catalogue will enable genomic epidemiologists and researchers to annotate genomes for insertion sequences more extensively and advance knowledge of how insertion sequences contribute to bacterial evolution in general and antimicrobial resistance spread across microbial lineages in particular. This will be useful for genomic surveillance, and for development of microbiome engineering strategies targeting inactivation or removal of important transposable elements carrying antimicrobial resistance genes.

## Introduction

Swapping genetic information between members of a microbial community, a mechanism referred to as horizontal gene transfer (HGT), is a key process in the microbiome. It allows for the spread of new genes and functionality throughout the community. The result of HGT can be acquisition of a new gene, duplication of an existing gene or even interruption of a current gene. The mechanisms that support HGT have been well described and involve the transfer of mobile genetic elements (MGEs). MGEs are best defined as broadly as possible, as any genetic element that can mediate its own transfer from one part of a genome to another or between different genomes. The most complex elements are conjugative plasmids and Integrative Conjugative Elements (ICEs) which can mediate their transfer between bacterial cells^1^. The simplest and most abundant MGEs are the insertion sequences which only contain enough genetic information for their own transposition. MGEs are best thought of as a continuum ranging from the relatively simple insertion sequences right up to conjugative elements and everything in between^2^. MGEs are crucially important in bacterial evolution as a result of the extensive diversity they generate, an aspect of this is their central role in the spread of antimicrobial resistance genes (ARGs) between microbial genomes^3^.

Insertion sequences are short transposable elements between 700-2,500 bp in length containing genes that code for the proteins involved in their own transposition they are found in both chromosomes, ICEs and plasmids^4^. Most insertion sequences contain one or sometimes two genes encoding transposases, the most ubiquitous genes in prokaryotic and eukaryotic genomes^5^. Insertion sequences and transposons (transposons are defined at genetic elements that can transpose from one part of the genome to another but carry sequences other than those involved in transposition, unlike insertion sequences which just encode the genetic information for their own translocation) can be broadly classified by the amino acids in their transposase, commonly DDE (aspartic acid, aspartic acid and glutamic acid), DEDD or HUH (two histidine residues separated by any large hydrophobic amino acid) motifs, and their mechanism of transposition (either conservative or replicative)^3^. Common DDE insertion sequences contain two inverted terminal repeats (ITRs) at each end of a 10-50 bp size DNA sequence that are reverse complement sequences of each other. Some insertion sequences are flanked by unique shorter direct repeat sequences, also known as target site duplications (TSDs), which are formed by the duplication of the insertion sequence target site upon insertion^4^. Unit transposons are a similar type of transposable element to insertion sequences containing a pair of ITRs but can also carry ARGs as well as transposases^3^. For simplicity, the abbreviation “IS” will be used hereafter to mean insertion sequence or unit transposon. ARGs can also be carried by composite transposons that are bounded by two copies of two different insertion sequences which can move together in a single unit^6^. A composite transposon can contain one or more passenger genes, such as ARGs, flanked by two insertion sequences and with two TSDs at both ends^3^.

Microbial genomes can be annotated for ISs by querying reference databases of known transposable elements, such ISfinder^7^, but these databases are small and do not represent many transposable elements in nature. As transposable elements are the most ubiquitous and abundant MGE, it is a continual effort to catalogue them all using common methods. Novel ISs containing ITRs can be detected using computational tools, such as EMBOSS^8^, that search for palindromic sequences representing ITRs^9^. However, transposable elements in isolated genomes that are assembled from short reads can be misassembled or incomplete, since assembly algorithms struggle to resolve repeated elements^10^. Additionally, ITR pairs are not typically exact reverse complements, and algorithms that only detect perfect palindromes may fail to identify many insertion sequences. Alternatively, novel ISs can be identified by manually searching for ITRs or flanking regions of interest (such as ARGs) using a genome browser, but this can be a difficult and tedious process. Alternatively, Hidden Markov Models (HMMs) have been used to identify transposases within these elements, include those without ITRs^9^. However, the presence of a transposase is not sufficient evidence for a transposition event to have occurred.

In this paper, we present a tool called Palidis (Palindromic Detection of Insertion Sequences) that finds ISs using an efficient maximal exact matching algorithm to identify ITRs across different genomic loci in reads sequenced from mixed microbial communities. These ISs can then be pooled and clustered to create a non-redundant catalogue of ISs. PaliDIS can also predict the origins of these ISs by querying them against ISfinder or a COmpact Bit-sliced Signature (COBS) index^11^ of 661,405 microbial genomes^12^. Here, we present the theory and implementation of this tool on 264 short read metagenomes to generate 879 unique ISs included in the first release of the Insertion Sequence Catalogue (ISC). Beyond this paper, Palidis will continue identifying ISs to expand ISC.

### Theory and Implementation

Palidis is implemented as a Nextflow pipeline with all dependency software packaged in one container image. The input file of Palidis is a tab-delimited manifest text file that contains information on the read file IDs, the file paths to the read fastq.gz files, sample ID and file paths to the assemblies. The output files are a FASTA file of ISs and accompanying tab-delimited file of information. The following steps are also illustrated in Figure 1.

**Figure 1.**
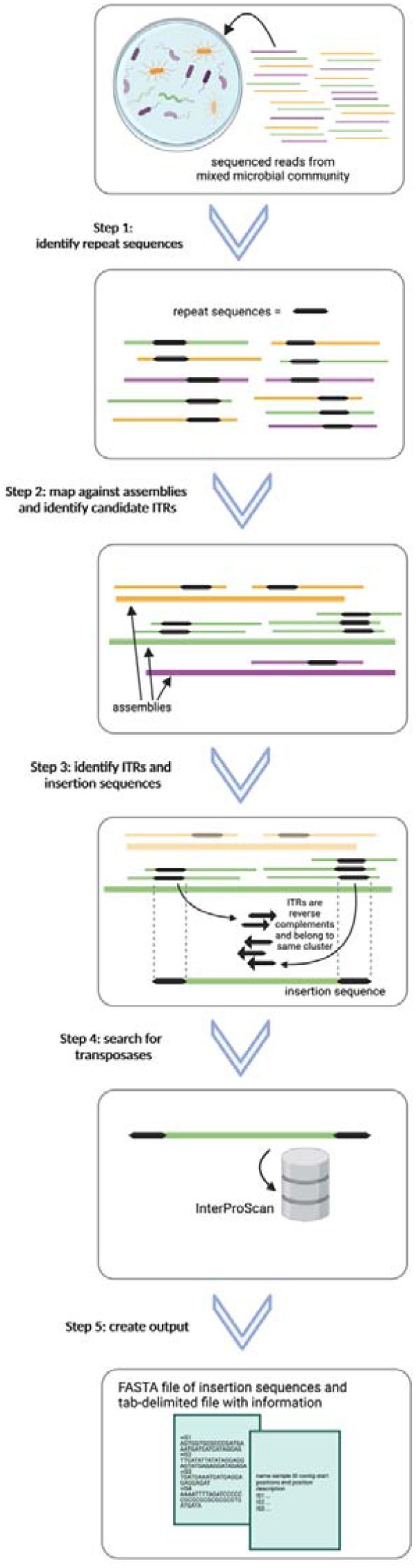
Steps summarising Palidis. Step 1: Reads from mixed microbial communities are pre-processed and run through pal-MEM to identify reads containing repeat sequences. Step 2: Reads containing repeat sequences are mapped against the assemblies using Bowtie2 to find their positions and proximity filters applied to identify candidate ITRs. Step 3: Candidate ITRs are clustered using CD-HIT-EST. ISs are identified by ITRs that are of the same cluster and are reverse complements of each other. Step 4: Search of transposases using InterProScan. Step 5: Final outputs of a FASTA file with insertion sequences and tab-delimited file with information are created.

### Step 1: Reads from mixed microbial communities are pre-processed and run through pal-MEM to identify reads containing repeat sequences

Firstly, the FASTQ files are converted to FASTA files with headers prepended with their sequence order (e.g. Seq1, Seq2 etc.). A software tool, called pal-MEM (https://github.com/blue-moon22/pal-mem), was developed and applied an efficient maximal exact matching algorithm^13^ to identify repeat sequences that may represent ITRs. A maximal exact match (MEM) between two strings is an exact match (i.e. an exact local alignment), which cannot be extended on either side without introducing a mismatch (or a gap).

### Preparing the reference and query data structures

pal-MEM creates a reference hash table from the sequences for some integer *k*>0 defined by the user, in which *k*-mers are the keys and the corresponding occurring positions are their values. The nucleotides of *k*-mers are encoded as unique combinations of two bits (0 and 1), (where A is 00, C is 01, G is 10 and T is 11), reducing memory requirements. In addition, it is not required for all *k*-mers to be stored, reducing the demand on memory further. A *k*-mer is stored only when it has a position that is a multiple of *(L – k) + 1* (where *k* is the length of the *k*-mer and *L* is the minimum ITR length), i.e.

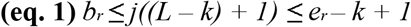

where *b*_*r*_ and *e*_*r*_ are the start and end positions of a maximal exact match (MEM) and *j* ≥ 1. The sequences are then also used to create a query data structure of unsigned 64-bit integers representing blocks of 32 nucleotides where each nucleotide is represented by two bits (A is 00, C is 01, G is 10 and T is 11). Random 20-bit sequences are stored between the array of reads define their boundaries. The start and end positions for each read and random sequence are stored in another data structure.

### Applying the algorithm to find repeat sequences

Each *k*-mer from the query read is looked up against the reference hash table to retrieve a matching *k*-mer. The first *k*-mer window starts from the beginning of the query and continues to shift every two bits, but skips the positions within the random sequences. These matching *k*-mers are then extended in both directions to make larger sequence matches until mismatches disrupt the extension, making a MEM. The algorithm performs this process using an interval halving approach. The sequence is extended to the left end position of the shortest of the two sequences. If there is no match, the extension is halved until a match is made. The extension is elongated by one nucleotide at a time until no more exact matches can be made. This is repeated on the right side. A repeat sequence is found once a MEM has a length greater than or equal to the minimum ITR length and less than or equal to the maximum ITR length as defined by the user. If a repeat sequence is found, pal-MEM will move on to the next read in the query, given it is expected that a read from short-read sequencing would contain only one ITR.

### Dealing with technical repeats from amplified read libraries

Read libraries are dominated by technical as well as biological repeated sequences that are the result of sequencing amplified regions. To reduce the frequency of technical repeats being identified as biological repeats, MEMs are also excluded if their start or end positions are within a buffer length of 20 nucleotides (40 bits) from either end of the read. This model represents an alignment of the prefix or suffix of a read typical of a technical repeat.

### Step 2: Reads containing repeat sequences are mapped against the assemblies using Bowtie2 to find their positions and proximity filters applied to identify candidate ITRs

Reads containing repeat sequences identified in Step 1 are mapped using Bowtie2^14^ against their associated assemblies. A Python script uses the output of Bowtie2 to identify mapped reads with candidate ITRs where the positions of the repeats are located between the minimum and maximum IS length as defined by the user.

### Step 3: Candidate ITRs are clustered using CD-HIT-EST. ISs are identified by ITRs that are of the same cluster and are reverse complements of each other

The candidate ITRs are clustered using CD-HIT-EST^15^ where nucleotide sequences that meet a 1) sequence identity threshold *c*, 2) a global *G* 1 or local alignment *G* 0, 3) alignment coverage for the longer sequence *aL*, 4) alignment coverage for the shorter sequence *aS* and 5) minimal alignment coverage control for the both sequences *A* (that can be specified by the user). The ISs are generated in a FASTA format with an accompanying tab-delimited file containing the sample ID, assembly name, start and end positions of the ITRs and their cluster using a Python script. The ISs must contain ITRs that 1) belong to the same cluster, 2) are within the minimum and maximum specified ITR length, 3) are within the minimum and maximum IS length, and 4) are reverse complements of each other where the two sequences aligned using BLASTn^16^ (with parameters *-task blastn -word_size 4*) have “*Strand=Plus/Minus*” and “*Identities”* greater than or equal to the specified minimum ITR length.

### Step 4: Search of transposases using InterProScan

Candidate ISs are queried for transposases using the InterProScan^17^, a tool that combines multiple search tools to predict protein family membership. Putative ISs must have Transposase, Integrase-like and/or Ribonuclease H within at least one protein family description.

### Step 5: Final output

A Python script generates a FASTA file of ISs and a tab-delimited file of information including: 1) their name (containing information on the length of the IS, the InterPro or PANTHER accession(s) identified in Step 4 and their position(s)), 2) sample ID, 3) contig, 4) start and end positions of the two ITRs on the corresponding contig, and 5) description of the protein family represented by the accession(s).

### Using Palidis to create Insertion Sequence Catalogue v1.0.0

A catalogue of insertion sequences was generated using Palidis applied to 264 human oral and gut metagenomic reads from the Human Microbiome Project (Supplementary Data 1)^18^. The reads were quality controlled, filtered and assembled as previously described^19^. A total of 2,517 ISs were identified from 1,837 contigs across 218 (out of 264) samples with Palidis v3.1.0 using default parameters (*--min_itr_length 25 --max_itr_length 50 --kmer_length 15 -- min_is_len 500 --max_is_len 3000 --cd_hit_G 0 --cd_hit_c 0*.*9 –cd_hit_G 0 -cd_hit_aL 0*.*0 -- cd_hit_aS 0*.*9*) (Supplementary Data 2).

The ISs were then clustered using CD-HIT-EST v4.8.1 (with a sequence identity threshold *–c 0*.*95* and default parameters) to create the Insertion Sequence Catalogue (ISC). ISC contains a FASTA file of 879 unique ISs between 524 and 2999 bp in length (Fig. 2a) and containing 87 unique transposases (Fig. 2b) (https://github.com/blue-moon22/ISC).

**Figure 2.**
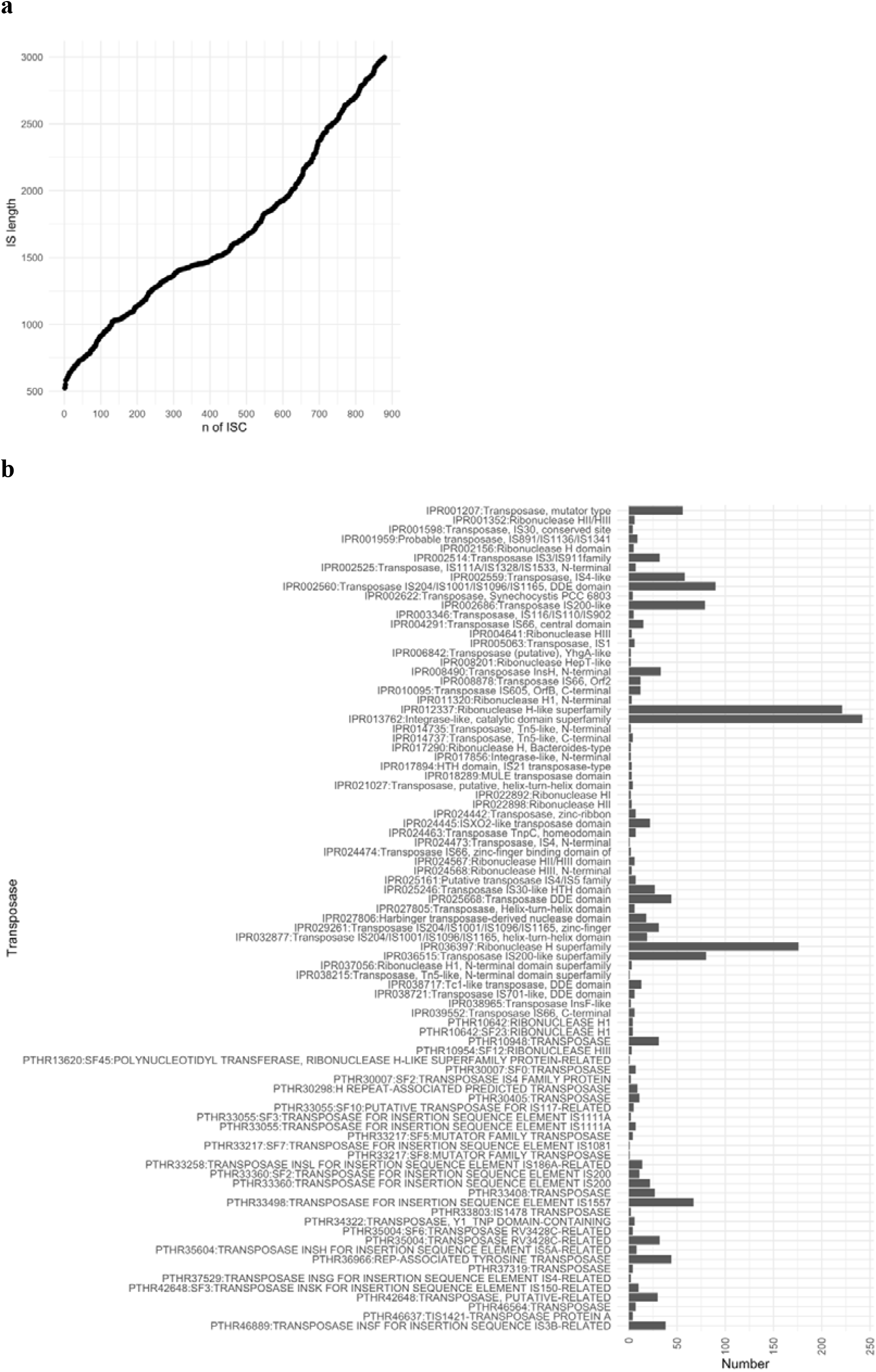
**a)** Length of insertion sequences and **b)** Number of transposases within the ISC

In order to identify ISs from ISC that have previously been discovered, ISC was queried against ISfinder using their online BLAST search tool (https://www-is.biotoul.fr/search.php) (with *e-value 0*.*01* and using default parameters on 7^th^ October 2022). 360 (41.0 %) ISs have hits in ISfinder, while the remaining 519 are novel. 60 have a strict homology with ISs from ISfinder (*e-value < 1e-50*), while the other 300 have a loose homology (*0*.*01 > e-value* ≥ *1e-50*).

The origins of the ISs were determined by querying the ISC against a COBS index of 661,405 bacterial genomes (referred to as the 661k database) from European Nucleotide Archive (ENA) (http://ftp.ebi.ac.uk/pub/databases/ENA2018-bacteria-661k/661k.cobs_compact)^12^ using cobs query v0.1.2 (with *-t 0*.*9* and using default parameters).

155 ISs were located in 791 samples (NCBI BioSample IDs) from the 661k database. Metadata was available for 785 for these samples from the 661k database study^12^. 684 samples came from isolates that had an associated taxonomic identity, whereas the other 85 samples were sequenced from microbial communities or were unclassified (known as “bacterium”). These 684 samples originate from 63 known genera (Fig. 3a) with 138 known species (not labelled *sp*.) (Fig. 3b).

**Figure 3.**
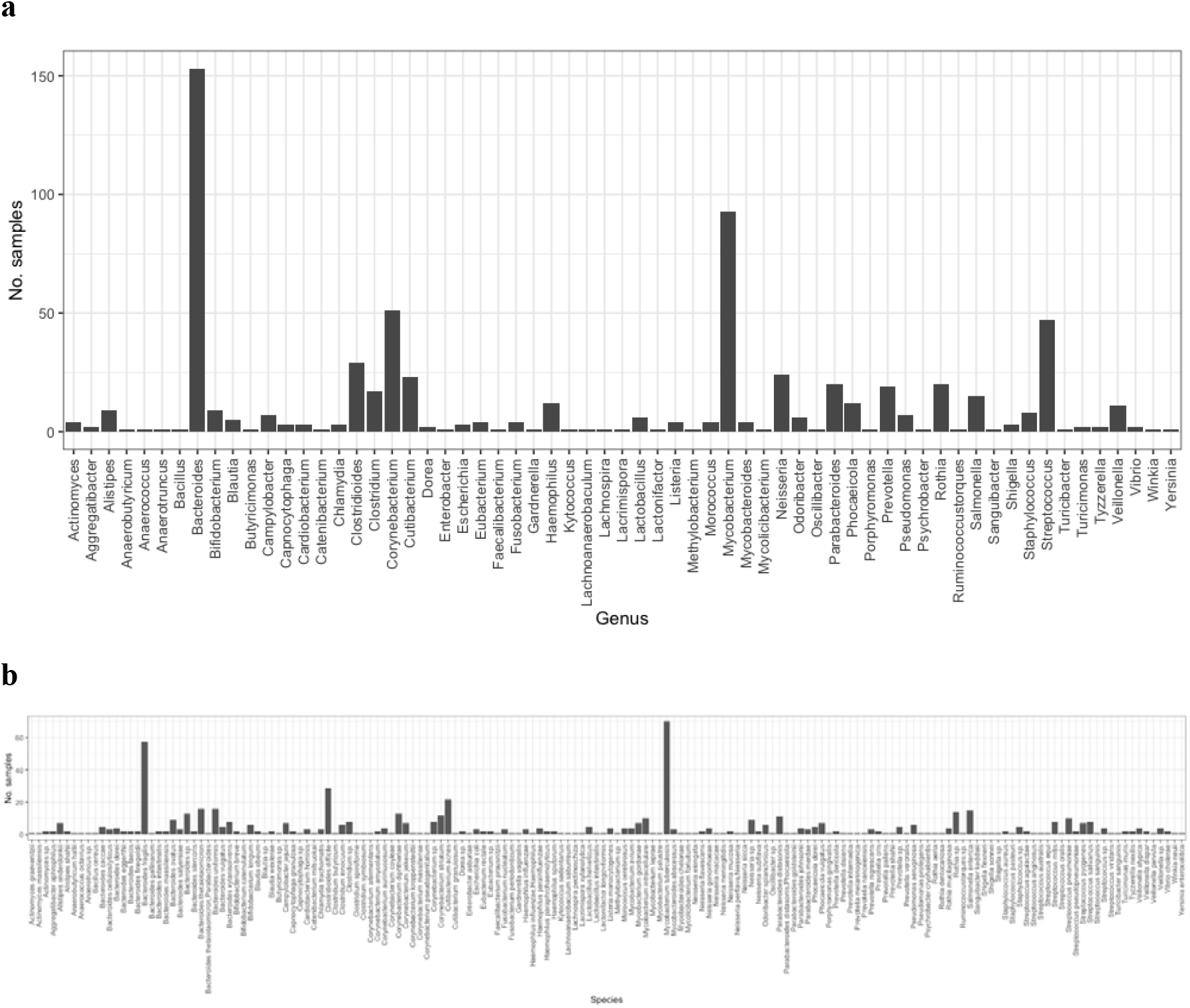
Number of samples from the 661k database that contain an IS found in across **a)** genera; **b)** species in the 661k database

Many ISs were shared across bacterial genomes of different species, genera and classes. 52 and 70 ISs originate from more than one known genus (Fig. 4a) and known species (not labelled *sp*.), respectively. The IS that is shared across most genera (21 genera and 46 species) and was also found in ISfinder as IS1249, is IS_length_1391-IPR001207_495_804 (Fig 4b, blue square), containing a Transposase, mutator type. The IS that is shared across the most genera but was not found in ISfinder, is IS_length_2555-IPR036397_1291_1768-PTHR35004_877_2065-IPR012337_1315_1720, containing a RV3428C-related Transposase (Fig 3b, red square). Many ISs found in multiple *Bacteroides, Corynebacterium, Curibacterium* and *Prevotella* species are solely represented by those not found in ISfinder, suggesting that in its current release (7^th^ October 2022), ISfinder database is underrepresenting ISs in these genera (Fig. 4c).

**Figure 4.**
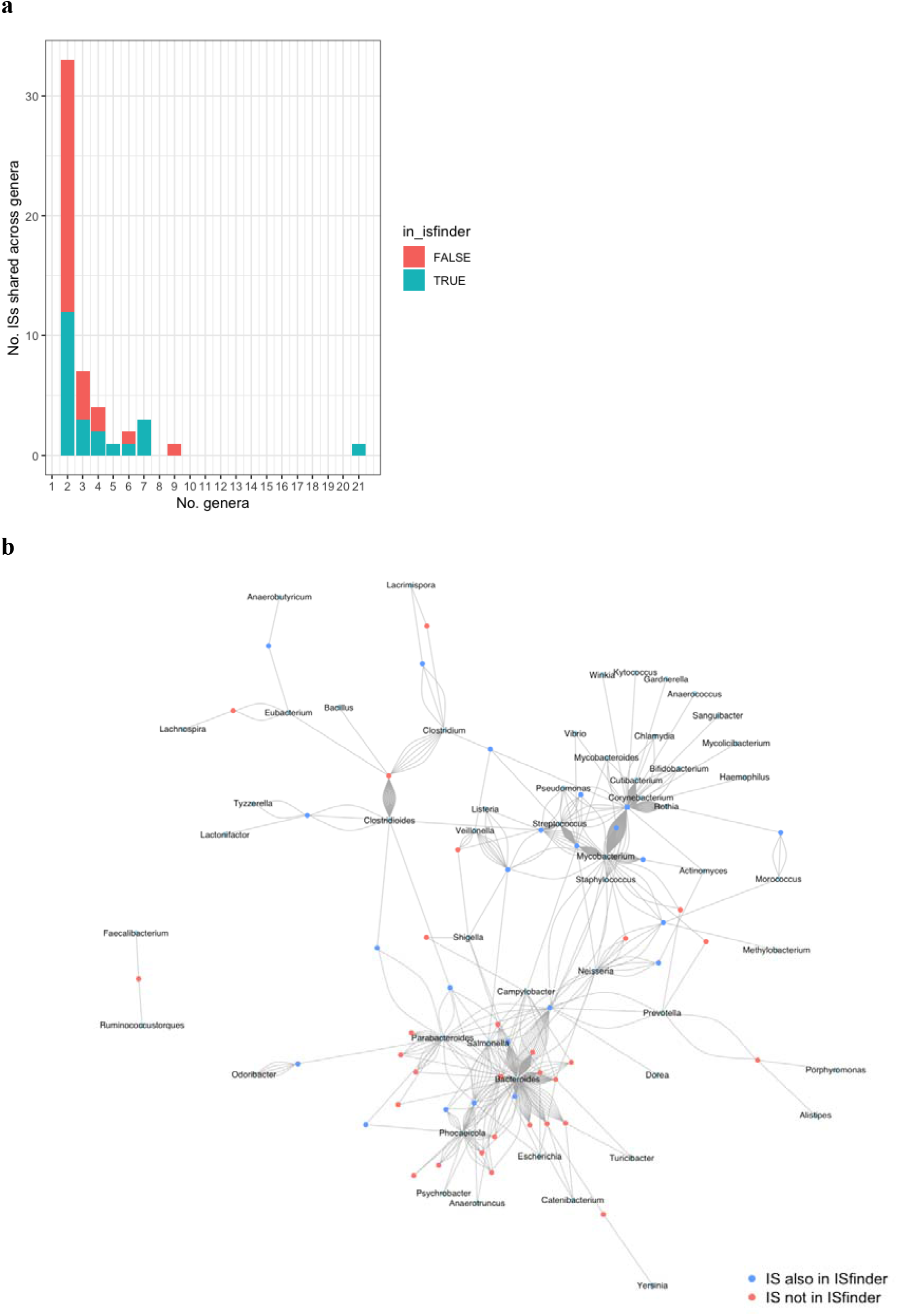
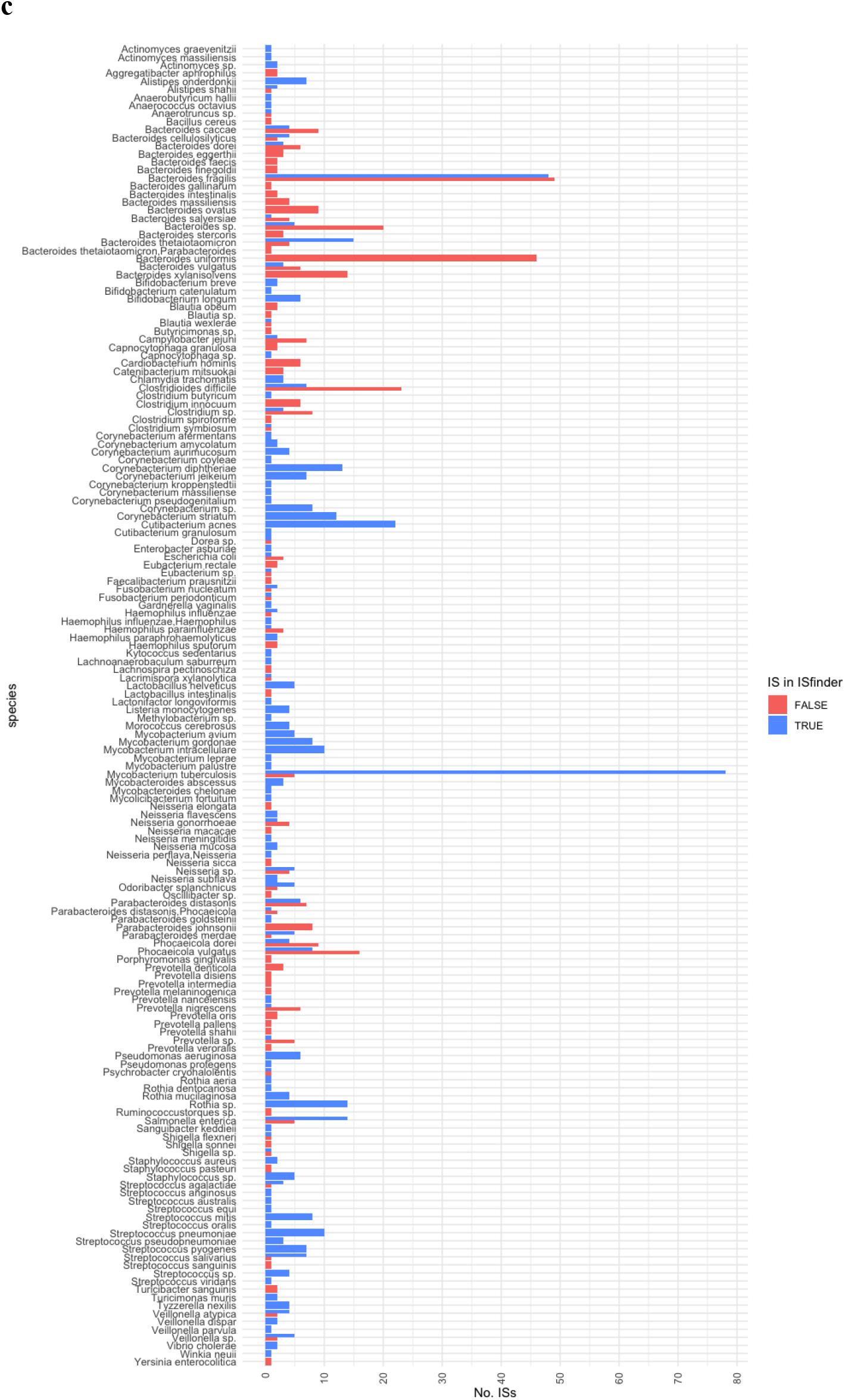
**a)** Number of ISs shared across a number of distinct, known genera in the 661k database; **b)** Network of shared ISs between genera. Each vertex represents either an IS (in ISfinder: blue or not in ISfinder: read) or a genus (labelled); **c)** Number of ISs that are either in ISfinder (blue) or not (red) that are in distinct, known species in the 661k database

## Discussion

Identification of transposable elements, including insertion sequences, in metagenomic datasets is critical in our ability to accurately define the profile of mobile genetic elements. In turn, accurate and complete characterisation of mobile genetic elements (i.e. the mobilome) of a community is central to understanding the spread and epidemiology of different genes in microbial communities, such as virulence genes and antimicrobial resistance genes. Here, we describe a tool and subsequent catalogue that enables this to proceed. Palidis is a tool that discovers novel ISs from mixed microbial communities by applying a fast maximal exact matching algorithm to identify ITRs. As a result, we have released the first version of ISC, a catalogue containing 879 ISs. Already, this is a valuable resource for researchers to search for ISs in isolated genomes. However, since Palidis was only applied to metagenomes sequenced from the healthy human oral cavity and stool samples, it is recommended ISC v1.0.0 is used as a reference for annotating isolates sourced from human oral and stool samples.

The main limitation of the current ISC is that it only contains common DDE types of ISs with ITRs, although these mobile genetic elements make up a large proportion of ISs^3^. Palidis is currently only equipped with discovering ISs with ITRs. We are planning to include other databases into the catalogue, such as ISfinder, and we invite the research community to contribute and submit ISs to the catalogue. Another limitation is that the catalogue currently contains ISs with ITRs that are 25 or greater nucleotides in length as generated by Palidis, although ITRs can be as short as 10 nucleotides in length. It is possible to run Palidis with a lower minimum ITR length threshold and smaller *k*-mer length, but at these smaller sizers, it becomes more computationally intensive, especially with more complex mixed microbial genomes. However, we will run Palidis with a lower minimum ITR length threshold on less complex genomes to discover ISs with smaller ITRs.

It is also important to note that all ISs in the catalogue contain a region that is flanked by ITRs within a 500 to 3000 bp proximity. Given the recursive mechanism of insertion events (i.e. ISs inserting within ISs), it is possible for a region to also contain another IS. Therefore, it is also possible for regions that have been lengthened by other insertion events to extend outside this proximity range and be missed by Palidis. Increasing the maximum IS length will account for this, and may be done for future iterations of the ISC.

In light of current times, disruptive sequencing technologies are advancing rapidly by becoming more accurate and generating longer reads. Very soon, Palidis could be applied to longer reads of isolates to identify novel ISs, as well as generating reference databases (like ISC) from mixed microbial genomes. However, the ISC could become a valuable resource for querying microbial genomes for ARGs that have been acquired through transposition. We will continue to enrich the ISC towards a comprehensive catalogue by applying Palidis with different parameters to more mixed microbial genomes from a diverse range of sources, and encouraging submission of ISs from the scientific community.

## Abbreviations

ARG: antimicrobial resistance gene
bp: base pairs
ISC: Insertion Sequence Catalogue
IS: insertion sequence/unit transposon
ITR: inverted terminal repeat
MEM: maximal exact match

## Acknowledgements

The project was supported by the Centre for Host-Microbiome Interactions, King’s College London, funded by the Biotechnology and Biological Sciences Research Council (BBSRC) grant BB/M009513/1 awarded to D.L.M. S.S. was supported by Engineering and Physical Sciences Research Council (EPSRC), EP/S001301/1, Biotechnology Biological Sciences Research Council (BBSRC) BB/S016899/1 and Science for Life Laboratory (SciLifeLab). S.P.P. is supported in part by the PANGAIA and ALPACA projects that have received funding from the European Union’s Horizon 2020 research and innovation programme under the Marie Sklodowska-Curie grant agreements No. 872539 and 956229, respectively.

